# Human Integrator provides a quality checkpoint during elongation to facilitate RNA polymerase II processivity

**DOI:** 10.1101/2023.02.17.528960

**Authors:** Sara Rohban, Mahmoud-Reza Rafiee, Jernej Ule, Nicholas M. Luscombe

## Abstract

Integrator is a multi-subunit complex that directly interacts with the C-terminal domain (CTD) of RNA polymerase II (RNAPII). Through its RNA endonuclease activity, Integrator is required for 3′-end processing of both non-coding and coding transcripts. Here we demonstrate that depleting Integrator subunit 11 (INTS11), the main catalytic subunit of the Integrator complex, leads to a global elongation defect as a result of decreased polymerase processivity. We observe this defect in the region approximately 12 to 35 kb downstream of the transcription start site (TSS), where RNAPII normally transitions to its maximum processivity. We also identify an important role for INTS11, possibly in association with RNAPII CTD phospho-Tyr1, in repressing antisense transcription upstream of active promoters, as well as repressing transcription of genic regions near AsiSI-induced double-strand breaks.

Altogether, this study points toward a novel function of Integrator in promoting termination of incompetent RNAPII molecules while facilitating the transition to fully processive polymerase in order to enable efficient elongation.

## Introduction

RNA polymerase II (RNAPII), the central enzyme of gene expression, synthesizes all messenger RNAs and the vast majority of small regulatory RNAs in eukaryotes. Precise execution of gene expression programs happens at steps of initiation, elongation and termination and is critical to cellular development and homeostasis and to the cellular response to different environmental cues. The highly repetitive carboxy-terminal domain (CTD) of RPB1, the largest subunit of RNAPII, plays a central role in the complex regulation of genes. CTD is composed of multiple tandem heptapeptides with the evolutionary conserved consensus motif Tyr1–Ser2– Pro3–Thr4–Ser5–Pro6–Ser7. RNAPII CTD can be phosphorylated at several sites and can serve as a docking platform for binding factors required for the regulation of the expression of RNAPII-transcribed genes(Phatnani and Greenleaf 2006). Among many interaction partners, the Integrator complex (INT) directly interacts with RNAPII CTD, possibly through its Tyr1-phosphorylated form(Baillat et al. 2005; Shah et al. 2018; Fianu et al. 2021). This multimeric protein complex is evolutionarily conserved in metazoans and contains at least 14 unique proteins known as core Integrator subunits (INTS1–INTS14). The pivotal subunit to Integrator function is Integrator subunit 11 (INTS11), which belongs to the metallo-β-lactamase superfamily and is a closely related homolog of CPSF-73, the endonuclease for pre-mRNA 3′-end processing. Together with INTS9 and INTS4, INTS11 forms a heterotrimeric complex called the Integrator cleavage complex that is responsible for Integrator’s endonucleolytic activity(Albrecht et al. 2018). INTS6, together with Protein Phosphatase 2A (PP2A), forms a distinct module with phosphatase activity, which associates with the Integrator core through interaction with INTS8(Huang et al. 2020; Zheng et al. 2020).

Initial characterization of the Integrator complex revealed its role in the 3′ end processing of RNAPII-mediated small nuclear RNAs (snRNAs)(Baillat et al. 2005). Further studies expanded the role of the Integrator complex to 3′-end cleavage of enhancer RNA (eRNA) primary transcripts(Lai et al. 2015) and human telomerase RNA (hTR)(Rubtsova et al. 2019), leading to transcriptional termination of these non-coding RNAs. Recently, at protein-coding genes, Integrator was shown to have a broader regulatory role at multiple stages during transcription, such as regulating transcription initiation and RNAPII pause release(Gardini et al. 2014; Huang et al. 2020), transcriptional attenuation(Elrod et al. 2019; Lykke-Andersen et al. 2021), transcriptional elongation(Beckedorff et al. 2020) and termination of transcription at diverse classes of gene targets(Skaar et al. 2015; Stein et al. 2022).

These findings suggest that Integrator plays a wide-ranging role in the regulation of gene expression. By applying the TT_chem_-seq method, which provides a high-resolution transcriptome profile of nascent RNAs, we found that depletion of the Integrator main catalytic subunit, INTS11, results in defective RNAPII elongation at coding genes, starting within a window of 12–35 kb from the TSS. We also observed aberrant expression of antisense RNA at promoters, and spurious transcription near double-strand break (DSB) sites upon INTS11 depletion. Therefore, our analyses add additional evidence on the role of the Integrator complex in the quality control of RNAPII molecules, ensuring proper and efficient elongation, while decreasing non-productive and spurious transcription.

## Results

### INTS11 knockdown causes preferential down-regulation of long genes

To study how depletion of the catalytic components of Integrator affects transcription, we depleted INTS9 or INTS11 by transfecting specific siRNAs (siINTS9 or siINTS11, respectively) into the human osteosarcoma U2OS cell line. As a control, we used an siRNA without any known target in the human genome (hereafter called “siCtrl”). Consistent with what others have shown(Beckedorff et al. 2020), we found that depletion of either INTS11 or INTS9 led to the reduction of both factors at the protein level but not at the mRNA level (Fig. 1a,b). These data supported the previous finding that the two catalytic subunits of the Integrator complex are tightly associated(Beckedorff et al. 2020).

**Figure 1.**
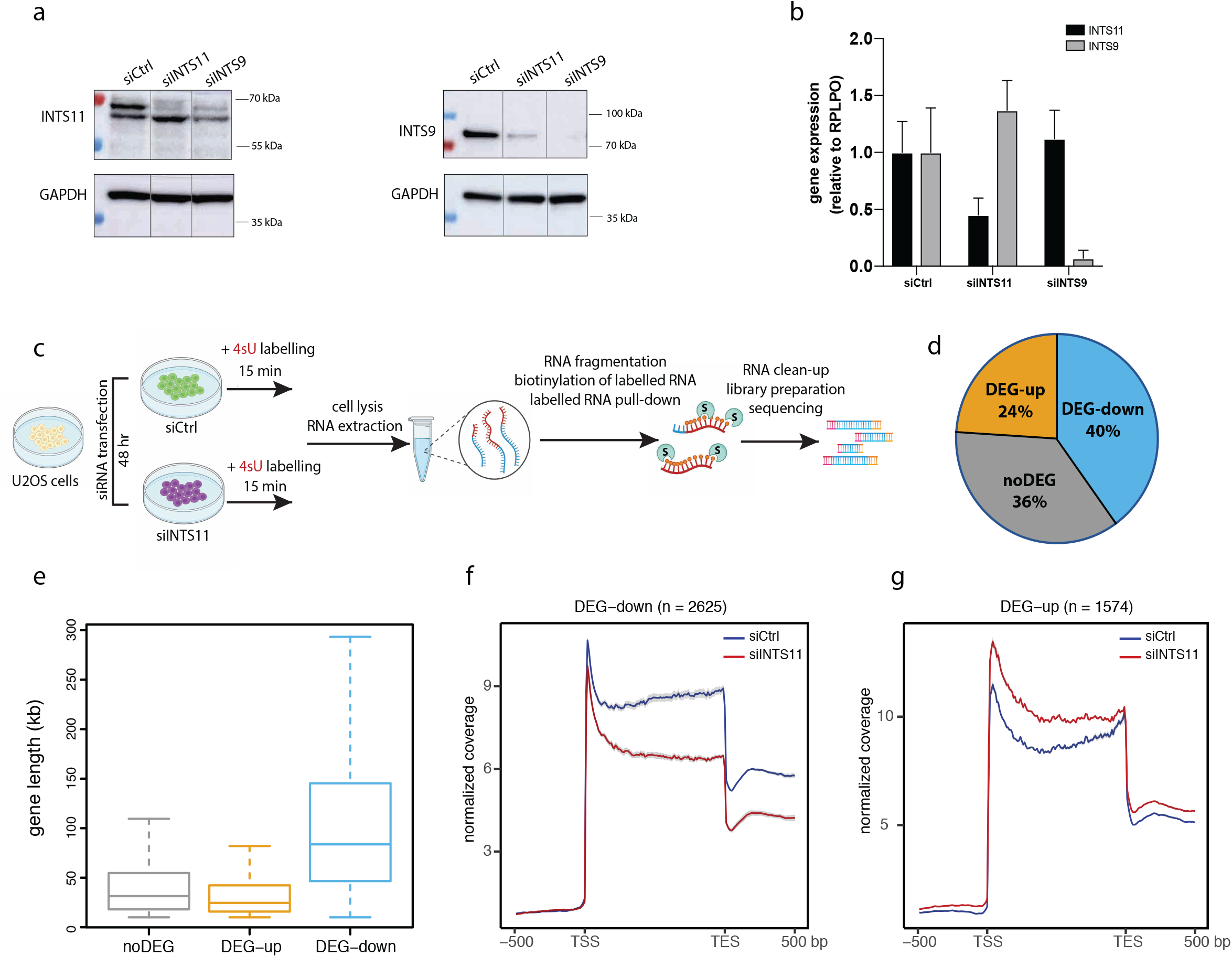
INTS11 knockdown in U2OS cells results in substantial transcriptional repression among protein-coding genes. (a) Western blot analysis showing INTS11 (upper left panel) and INTS9 (upper right panel) in U2OS cells transfected with siRNAs against INTS9 and INTS11 at 48h. Cells transfected with a non-targeting siRNA (siCtrl) act as the negative control. GAPDH was used as a loading control. (b) Quantitative RT-PCR analysis of INTS11 (black bars) and INTS9 (grey bars) expression levels in cells transfected with siCtrl, siINTS11 and siINTS9 after 48h. mRNA expression level was normalized with a housekeeper gene RPLPO and is plotted relative to control cells. (c) Experimental design of the TT-seq experiment on U2OS cells transfected with siCtrl or siINTS11 after 48h. (d) Pie chart of the differentially expressed genes upon INTS11 knockdown obtained from TT-seq data. Only active genes (RPKM>0.5) are included in the analysis. Up-regulated (DEG-up) and down-regulated (DEG-down) genes were defined as (p-adj < 0.05 and log2 fold change > 0) and (p-adj < 0.05 and log2 fold change < 0) respectively. (e) box plot showing gene length (kb) for the noDEG, DEG-up and DEG-down gene categories. (f and g) Metagene plot of TT-seq coverage in the DEG-down (e) and DEG-up (f) categories. TSS and TES denote transcription start site and termination site respectively.

For the rest of this work, we therefore chose to perform all experiments by depleting the main catalytic subunit INTS11. We first studied the genome-wide effect of this depletion on the dynamics of transcription. We used the transient transcriptome sequencing method TT_chem_-seq(Gregersen et al. 2020) to collect and sequence nascent RNAs in siCtrl- and siINTS11-transfected cells. In brief, newly synthesized transcripts were labelled by incubating cells in a medium containing 4-thiouridine (4sU) for 15 min. Following a fragmentation step by sodium hydroxide, the 4sU-labelled RNAs were biotinylated, captured by streptavidin and, after library preparation, subjected to sequencing (Fig. 1c).

At first, we sought to identify gene expression changes in the INTS11-knockdown cells compared with control cells. By applying the DESeq2 software(Love et al. 2014) to the TT-seq read counts on all protein-coding genes > 10 kb, we identified ~ 65 % (4199 out of 6464 genes) of the active genes (RPKM > 0.5 across all samples) as being differentially expressed. Interestingly, ~ 63 % [2625 genes] of these differentially expressed genes (DEGs) were down-regulated (DEG-down), and only ~ 37 % [1574 genes] were up-regulated (DEG-up), suggesting substantial transcriptional repression occurring in INTS11-depleted cells (Fig. 1d).

To obtain more insights into the cause of the transcriptional repression observed in these cells, we compared gene lengths in the DEG-up and DEG-down categories. We found that the down-regulated genes are generally longer than the up-regulated genes and those genes whose expression was not affected upon INTS11 knockdown (Fig. 1e). Two other siRNAs targeting other parts of the INTS11 mRNA also gave the same results (Fig. S1), indicating that the effect is unlikely to result from off-targeting.

Next, we looked at the metagene profile of the down-regulated genes, which mostly consist of long genes. Metagene analysis of the DEG-down cluster showed a drastic reduction in the 4sU signal in the gene body and toward the 3′ end of the genes in the siINTS11-transfected cells compared with control cells (Fig. 1f). Interestingly, the progressive reduction in the signal toward the 3′ end of the genes was also evident in the meta-profile of the DEG-up cluster (Fig. 1g), suggesting a global elongation defect in INTS11-depleted cells.

### INTS11-depleted cells exhibit an elongation defect

To investigate the effect of INTS11 knockdown on transcription elongation, we divided the protein-coding genes into 3 classes based on their gene length: short (10–20 kb), medium (20–50 kb) and long (> 50 kb). Average metagene analysis of TT-seq signal on short genes (< 20 kb) revealed an increase in the read density within gene bodies in INTS11 knockdown condition (Fig. 2a). This is consistence with the recent study by Stein et al., showing a significant increase in PRO-seq signal in early gene body transcription upon INTS11 degradation, leading to upregulation of short genes(Stein et al. 2022).

**Figure 2.**
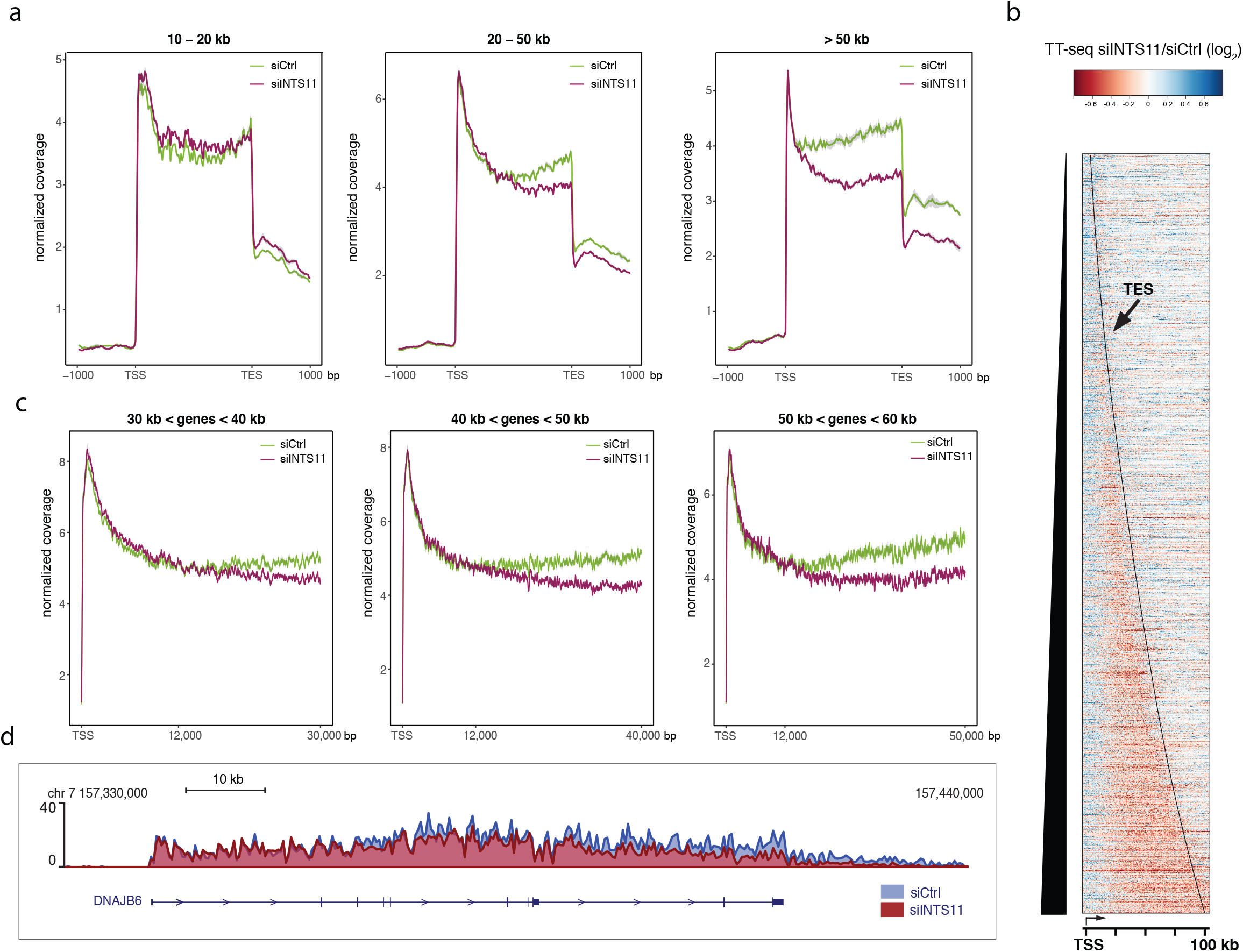
INTS11-depleted cells exhibit a global elongation defect. (a) Average gene body profiles of TT-seq signal for expressed protein-coding genes in three classes of genes based on their length (short 10-20 kb, medium 20-50 kb and long > 50 kb). (b) Normalized siINTS11/siCtrl ratio heatmap across the entire length of all expressed genes up to 100 kb, ordered by gene length. Only genes on the sense strand are plotted. The line in the plot shows the termination site (TES). (c) TT-seq signal density at three different classes of genes (divided by their gene length) in distance to the TSS along the x-axis. (d) An example of the TT-seq track (nascent RNA expression) at the *DNAJB6* locus in cells transfected with siCtrl and siINTS11, overlaid in blue and red respectively.

Moreover, our metagene analysis on medium and long genes demonstrated a significant correlation between the gene length and the 4sU signal, with the long genes showing the greatest signal reduction (Fig. 2a). To investigate whether the observed elongation defect is gene length dependent or not, we decided to visualize the profiles without gene length normalization. For this purpose, we generated heatmaps of siINTS11/siCtrl ratio with fixed binning, for genes up to 100 kb in length. As shown in Fig. 2b, TT-seq density was not significantly changed around the TSS of most of the genes upon INTS11 depletion. However, a remarkable global decrease in the signal was evident several kilobases downstream of the TSS into gene bodies in these cells. To pinpoint more precisely where the reduction in read density happens, we looked at the metagene profile of the genes downstream of their TSSs. This analysis revealed that the drop in 4sU signal in INTS11-depleted cells took place approximately 12 kb from the TSS in all genes regardless of their gene length category (Fig. 2c). This is the region where RNAPII elongation was shown to transition into a gradual acceleration phase to reach a final state of maximum elongation speed(Danko et al. 2013), thus indicating that INTS11-knockdown cells have a defect in transitioning into maximum elongation within genes longer than 15 kb.

### The elongation defect in INTS11-depleted cells results from loss of RNA polymerase processivity

Transcriptional output across the gene body is determined by two factors: (i) elongation rate, defined as the number of nucleotides transcribed per minute and (ii) processivity, defined as the number of nucleotides transcribed per RNAPII molecule. We, therefore, aimed to investigate whether the observed decrease in RNA synthesis in INTS11-knockdown cells results from reduced elongation rate or loss of RNAPII processivity. First, we calculated the siINTS11/siCtrl ratio of all expressed genes longer than 70 kb. This analysis revealed a linear decline of the TT-seq signal starting from TSS + 12 kb up to TSS + 35 kb. Beyond this point, transcription constantly remained low, and there was no further loss of signal in the knockdown cells (Fig. 3a,b). This finding, however, did not provide enough evidence to support one or the other possibility regarding the cause of the elongation defect in INTS11-depleted cells. To determine whether INTS11 depletion affects the speed of RNAPII elongation rate, we performed DRB/TT-seq in the control and INTS11-depleted cells. Transcription is halted at the pause–release step by treating cells with 5,6-dicloro-1-beta-D-ribofuranosylbenzimidazole (DRB) for 3 h. After DRB washout, transcription is restarted and 4sU is added to the medium during the last 10 min before cell harvesting to track the kinetics of nascent RNA synthesis. We sequenced 4sU-labelled transcripts after 10, 20 and 30 min of DRB washout. In control cells, the transcription wave progressed with a typical elongation rate (~ 2 kb/min(Baluapuri et al. 2019; Narain et al. 2021)) as shown by metagene and single-gene analyses (Fig. 3c-f). Moreover, we did not observe any significant change in elongation speed in INTS11-knockdown cells compared with control cells, confirming that the elongation rate is unaffected in these cells.

**Figure 3.**
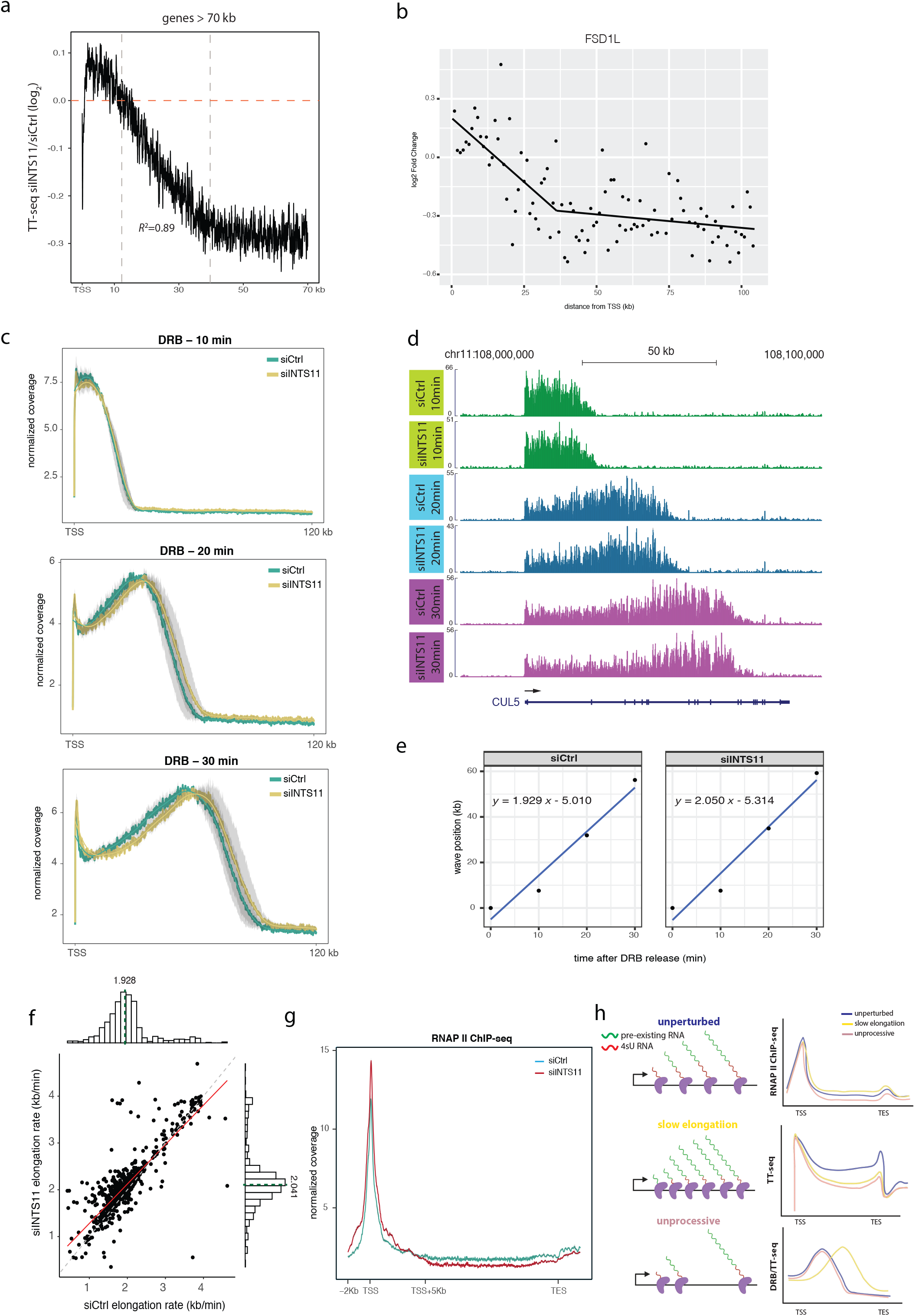
Transcription elongation defect in INTS11-depleted cells results from loss of RNAPII processivity. (a) Meta-profile showing siINTS11/siCtrl ratio of all expressed genes > 70 kb in the first 70 kb following the TSS. The dotted vertical lines demonstrate the TSS + 12 kb to TSS + 40 kb region. (b) Plot showing changes in nascent transcription upon INTS11 knockdown of an example gene (*FSD1L*). log2 fold change of TT-seq read counts on *FSD1L* gene in non-overlapping 1 kb windows is plotted. (c) Average metagene plots of DRB/TT-seq, showing the distribution of read density over the gene body from TSS to 120 kb downstream. Only genes > 120 kb were considered in the analysis. The fitted splines are also shown. (d) Example of DRB/TT-seq tracks of siCtrl- and siINTS11-transfected cells at 10, 20 and 30 min DRB release at the *CUL5* locus. (e) Calculation of RNAPII elongation rate in siCtrl- and siINTS11-transfected cells based on DRB metagene profiles at 10, 20 and 30 min post DRB release. Blue line shows linear regression through the data. (f) Scatterplot comparing elongation rates of 710 genes (gene length > 100 kb) between control and INTS11 knockdown-cells. Histograms with median values (dashed lines) are also shown. (g) Metagene plot from total RNAPII ChIP-seq. Normalized read coverage over all expressed genes > 50 kb is shown. (h) Left: schematic representation of RNAPII transcribing in an unperturbed, slow elongation and unprocessive manner. Right: changes in RNAPII ChIP-seq, TT-seq and DRB/TT-seq signal at each condition are compared.

We next performed chromatin immunoprecipitation sequencing (ChIP-seq) of total RNAPII in the siRNA-transfected cells. ChIP-seq of RNAPII gave a typical profile of RNAPII occupancy in the control cells (sharp peak proximal to TSS and low density on gene body). In INTS11-depleted cells, the read density also peaked around the pause site (the TSS); however, compared with control cells, RNAPII occupancy was reduced over the gene bodies (Fig. 3g). Altogether, the results from TT-seq, DRB/TT-seq and RNAPII ChIP-seq analyses support the idea that the elongation impairment we observed in INTS11-knockdown cells was due to the defects in elongation processivity rather than changes in elongation speed (Fig. 3h).

To rule out the possibility that the impaired elongation phenotype in INTS11-depleted cells was due to the reduced expression of important elongation factors, we calculated gene expression levels from total RNA-seq performed on siCtrl- and siINTS11-transfected cells. We did not find significantly altered expression of the components of major elongation factors such as the super elongation complex (SEC), or the PAF1 or FACT complexes (Fig. S2a), suggesting that the elongation defect in INTS11-depleted cells was not because of misregulation of key regulators of elongation. Furthermore, global splicing defects were not evident upon INTS11 depletion as determined by 5′ splice junction usage in siCtrl and siINTS11 cells (Fig. S2b), which argues against major changes in splicing efficiency upon INTS11 knockdown. Since Integrator has been implicated in snRNA biogenesis, we also measured the expression of U1 snRNA to investigate whether depletion of INTS11 affects the level of UsnRNAs. Northern blot analysis of U1 snRNA indicated that, under our experimental conditions, the level of mature U1 was unchanged in INTS11-depleted cells (Fig. S2c).

### INTS11 knockdown increases the abundance of promoter upstream antisense transcripts

A close inspection of the nascent RNA tracks in siCtrl- and siINTS11-transfected cells revealed that INTS11-depleted cells generally have higher levels of divergent antisense transcripts in the TSS-proximal genic regions (Fig. 4a). To consolidate this observation at the genome-wide level, we first calculated TT-seq signal density in sense and antisense directions of the TSS of all non-overlapping protein-coding genes. The average metagene analysis demonstrated a clear increase in the promoter upstream antisense transcription in INTS11-depleted cells compared with control cells (Fig. 4b). This phenotype could be further confirmed by computing the “antisense score” which was defined as the ratio (log_10_) of TT-seq signal in the 1.5 kb window upstream of non-overlapping protein-coding genes in the antisense direction over the first 1.5 kb of genes (in the sense direction). Antisense scores for siCtrl- and siINTS11-transfected cells indicated that the loss of INTS11 results in accumulation of antisense transcripts (Fig. 4c).

**Figure 4.**
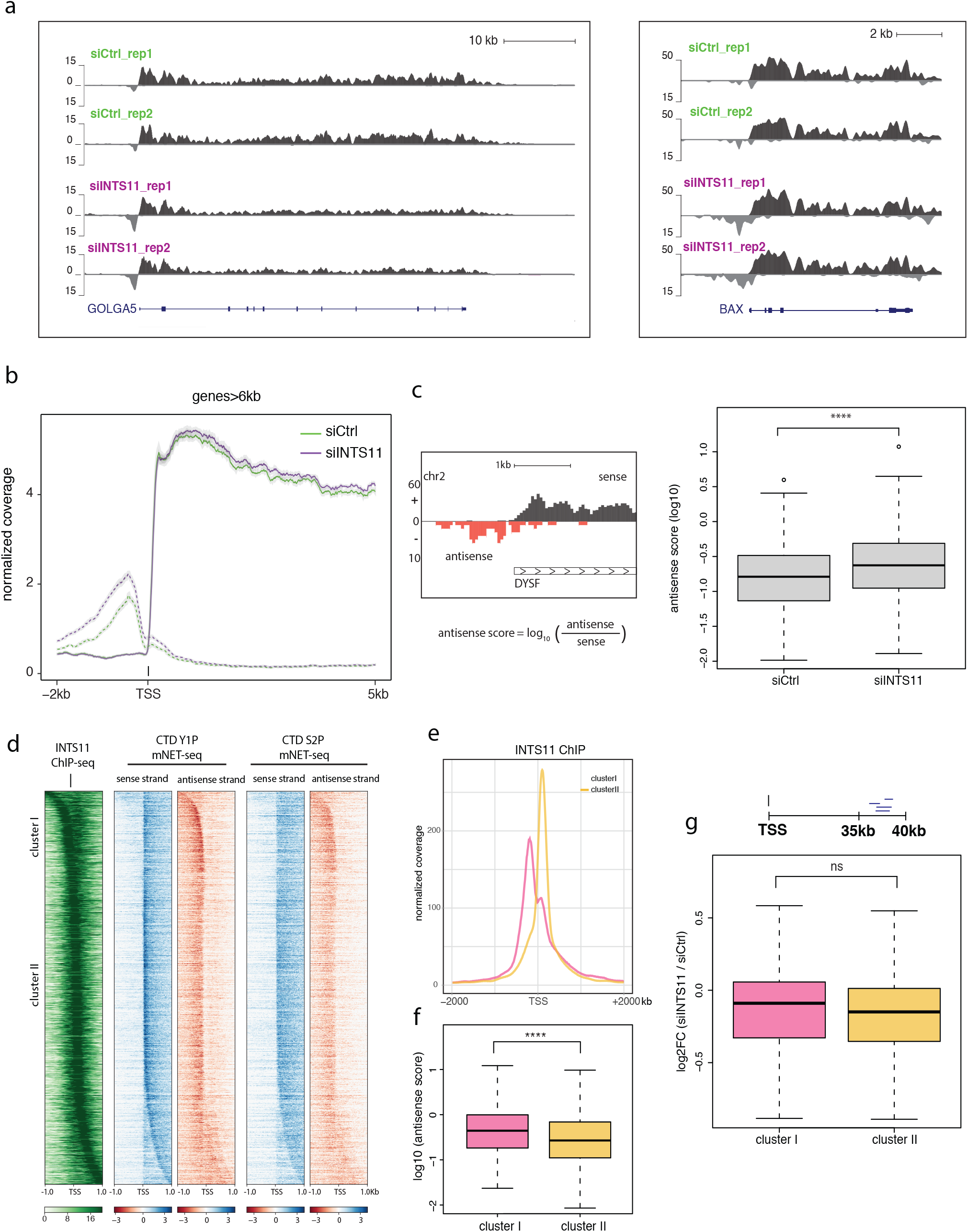
INTS11-depleted cells show a higher level of divergent antisense transcription. (a) Nascent TT-seq tracks in siCtrl- and siINTS11-transfected cells (replicate 1 and 2) for two example genes: *GOLGA5* and *BAX*. Sense transcription and antisense transcription are shown in black and grey respectively. Tracks are normalized by size factor and smoothened. (b) Average metagene profile of TT-seq signal in sense (solid line) and antisense (dashed line) orientation. 5 kb downstream and 2 kb upstream of TSS are shown. (c) Boxplot showing antisense score (log10) in siCtrl- and siINTS11-transfected cells. The nascent TT-seq signal on TSS - 1000 bp to TSS + 500 bp on antisense direction and the signal on TSS to TSS + 1500 bp on sense strand of the non-overlapping coding genes was taken. The ratio of antisense over sense signals (log10) was calculated as the antisense score, as depicted on the left (*t*-test p-value < 0.0001). (d) Heatmap representations of INTS11 ChIP-seq, CTD Y1P mNET-seq and CTD S2P mNET-seq signals aligned around TSS ± 1 kb. Data are ranked by INTS11 distance to TSS. Two clusters were defined based on the INTS11 binding distance relative to the TSS: cluster I with peaks within the window of TSS – 500 bp to TSS −100 bp and cluster II with peaks in TSS + 50 bp to TSS + 200 bp window. Blue and red represent mNET-seq signal on sense and antisense strands respectively. Only genes on sense strand are depicted. (e) Average distribution of INTS11 ChIP-seq signal aligned around TSS and divided into cluster I and II. (f) Box plot showing antisense score (log 10) in each cluster (*t*-test p-value < 0.0001). (g) Boxplot showing log2 fold change of sense transcription in cluster I and II. A window of TSS + 35 kb to TSS + 40 kb was used to assess the sense transcription. Only genes > 40 kb in each cluster are considered (*t*-test p-value = 0.233). Schematic representation of the analysis is shown above the plot.

Previous studies have identified RNAPII CTD phospho-Tyr1 (CTD 1YP) as a hallmark of antisense promoter transcription, suggesting its role in regulating the directionality of transcription(Descostes et al. 2014; Hsin et al. 2014). In line with these findings, using INTS11 ChIP-seq and CTD 1YP mNET-seq datasets, we observed co-enrichment of INTS11 and CTD 1YP near the promoters of genes (Fig. 4d). We then stratified genes based on the INTS11 binding distance relative to the TSS. We found that genes with a strong enrichment of INTS11 upstream of their TSS (TSS − 500 bp to TSS − 100 bp, cluster I) have significantly higher levels of CTD 1YP and antisense transcription than genes with INTS11 binding at their promoter regions (TSS + 50 bp to TSS + 200 bp, cluster II) (Fig. 4e,f). These data support a genome-wide effect of INTS11 on removing promoter upstream antisense transcripts and propose a model wherein Integrator in association with CTD 1YP modulates the expression of non-productive antisense transcription, thereby regulating the directionality of transcription. We next asked whether the increase in antisense transcription contributes to the elongation phenotype in INTS11-knockdown cells. To address this question, we compared the expression level of genes in each cluster (cluster I and cluster II) within a 5 kb window from TSS + 35 kb to TSS + 40 kb, where we observed a linear decline in the TT-seq signal. As shown in Fig. 4g, we did not find a significant difference in the expression level of genes between cluster I (with higher antisense score) and cluster II (with lower antisense score). Whether antisense and sense transcription are regulated independently, or whether they are linked together with regard to their dependence on INTS11, requires more investigation.

### Integrator is recruited to DSBs in an RNAPII-dependent manner to attenuate spurious transcription

We have shown that INTS11 has a crucial role in regulating RNAPII processivity. We conceived that Integrator-mediated termination might help to remove incompetent RNAPII complexes within 12–35 kb from TSS, thereby facilitating efficient transcription. We, therefore, wondered whether Integrator might also be important when transcription elongation is affected by an obstacle.

To answer this question, we took advantage of the AsiSI–ER cellular model to induce DSBs in transcriptionally active genes(Iacovoni et al. 2010). AsiSI is an 8 bp cutter restriction enzyme with a thousand annotated recognition sites in the human genome. Upon treatment with 4-hydroxytamoxifen (OHT), AsiSI–ER fusion proteins translocate into the nuclei of the overexpressing cells and generate site-selective DSBs across the genome. We observed that a subunit of the Integrator complex, INTS12, forms foci in response to OHT treatment in the U2OS cell lines harbouring the AsiSI–ER construct (Fig. 5a). Moreover, the INTS12 foci were co-localized with phospho-H2AX (γH2AX) foci, which mark DSB sites, confirming that INTS12 was indeed recruited to the sites of DNA damage in these cells (Fig. 5a,b). Furthermore, we found that the recruitment of INTS12 to the sites of damage was coordinated with the appearance of γH2AX foci upon DSB induction (Fig. S3a).

**Figure 5.**
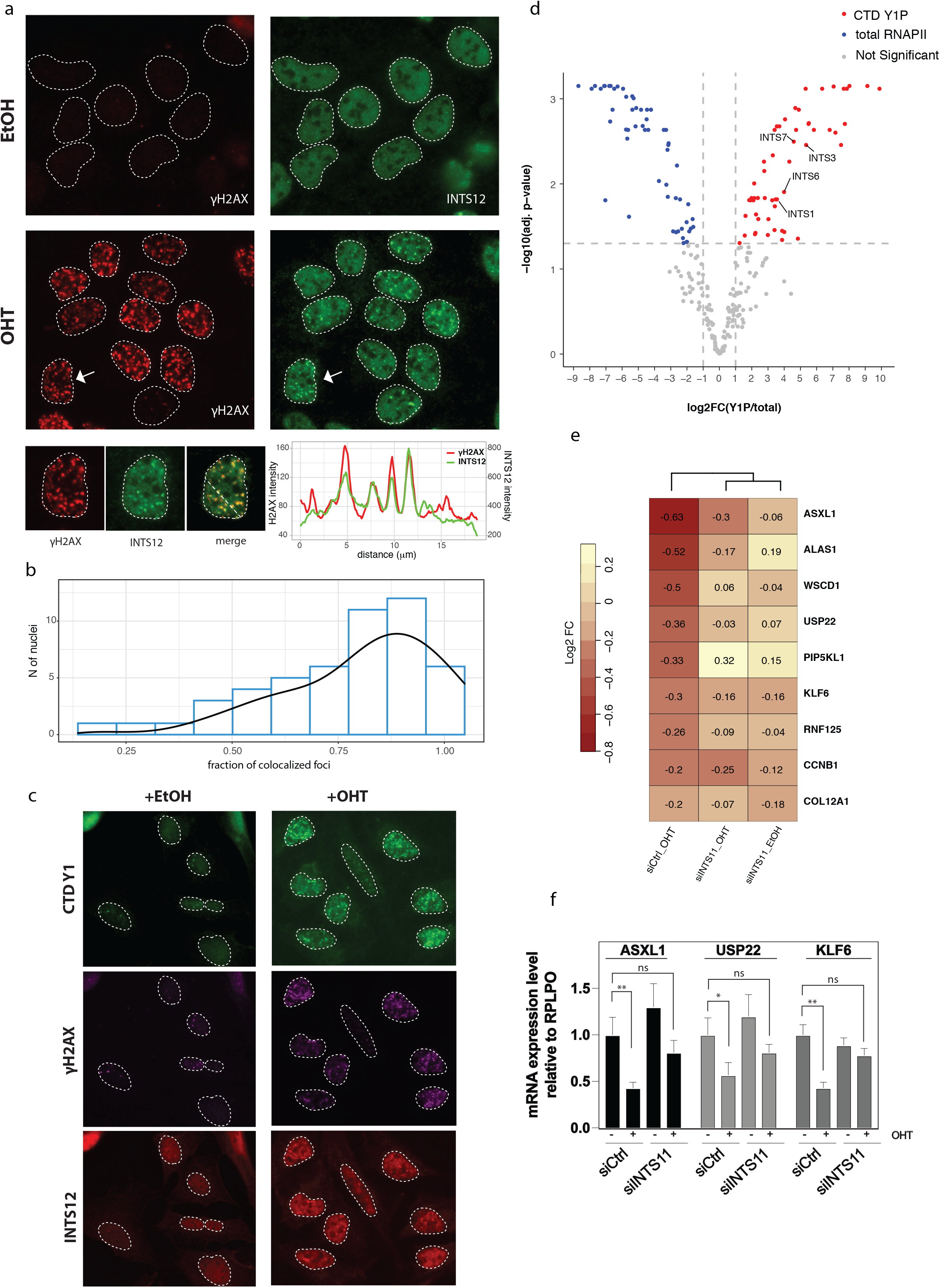
Integrator is recruited to the AsiSI-induced DSBs and is involved in transcriptional gene silencing upon formation of DSBs. (a) Immunofluorescence staining of yH2AX and INTS12 on AsiSI–ER–U2OS cells treated with OHT (to induce DSB) or ethanol (EtOH) as a control. An example of a merged nucleus (arrow) is shown below the panel. The RGB profile on the right shows yH2AX and INTS12 co-localization. (b) Histogram showing the fraction of co-localized yH2AX and INTS12 foci from 50 nuclei. (c) Immunofluorescence analysis of CTD Y1P, yH2AX and INTS12 as in (a). (d) Mass spectrometry analysis of the proteins interacting with CTD Y1P vs total RNAPII on chromatin. Volcano plot shows proteins that are enriched (red) or depleted (blue) in CTD Y1P compared to total RNAPII interactome in AsiSI–ER–U2OS cells treated with OHT. (e) Heatmap of expression level of AsiSI genes based on DESeq2 log2 fold change (log2 FC) obtained from TT-seq data. A window of TSS to TSS + 9 kb was selected for gene expression analysis. log2 FC is reported relative to control cells (siCtrl). (f) Quantitative RT-PCR analysis of expression of three selected AsiSI target genes in AsiSI–ER–U2OS cells transfected with siCtrl or siINTS11 and treated with OHT or ethanol (EtOH). mRNA expression level was normalized with a housekeeper gene RPLPO and is plotted relative to control cells (siCtrl + EtOH). Primers are designed to anneal to intron 1 of the genes. Data are presented as mean ± standard deviation (two-way ANOVA, * P < 0.05, ** P < 0.01, ns not significant).

To gain insight into the possible role of INTS12 at DSB sites, we set out to understand if the recruitment of INTS12 to these sites is dependent on RNAPII. We first pre-treated the AsiSI–ER–U2OS cells with the transcription inhibitors ⍺-amanitin or triptolide for 3 h before inducing DSBs. Immunofluorescence staining confirmed the occurrence of γH2AX foci in OHT-treated cells that were pre-treated with the transcription inhibitors, indicating that the transcription inhibitors ⍺-amanitin and triptolide did not affect the OHT-induced γH2AX signal. By contrast, the transcription inhibitors suppressed the formation of INTS12 foci upon OHT treatment (Fig. S3b), suggesting that INTS12 was recruited to the DSB sites in an RNAPII-dependent manner.

Recent findings on the involvement of RNAPII at DNA damage sites showed formation of RNAPII foci, predominantly CTD Y1P, at DSBs, which play an important role in recruitment of key DNA damage response factors to the damage sites(Burger et al. 2019). In line with these findings, using immunofluorescence assay, we demonstrated that upon induction of DSBs in AsiSI–ER cells, CTD Y1P form foci that are co-localized with γH2AX foci. Furthermore, we found that INTS12 foci co-localized with CTD Y1P and γH2AX in these cells (Fig. 5c). This observation is in agreement with our mass spectrometry data using CTD Y1P ChIP-SICAP (ChIP combined with selective isolation of chromatin-associated proteins(Rafiee et al. 2016)), showing that several subunits of the Integrator complex were associated with CTD Y1P in AsiSI–ER cells upon treatment with OHT. Importantly, Integrator co-localization with RNAPII complex is significantly higher using CTD Y1P in comparison to the pan-RNAPII antibody, suggesting the preferential association of the Integrator complex with CTD Y1P at the damage sites (Fig. 5d, supplementary table 1).

We then asked whether Integrator recruitment to the DSBs, mediated by RNAPII, is important for the reported DSB-induced gene silencing(Iannelli et al. 2017). To answer this question, we performed TT-seq on the AsiSI–ER–U2OS cells transfected with siCtrl or siINTS11 and treated with OHT or ethanol. We then looked at the expression level of genes that have an AsiSI cut site in the control and knockdown cells. Although around 1,200 AsiSI restriction sites have been mapped in silico in the human genome, not every site is efficiently cut by AsiSI. We took a published list of 74 genes located in proximity to the AsiSI sites (± 2 kb) that were detected by both BLISS (a method for breaks labelling in situ and sequencing) and γH2AX ChIP-seq(Iannelli et al. 2017).

This gene list was further filtered to include only genes whose expression (i) is significantly down-regulated upon OHT treatment in siCtrl-transfected cells and (ii) is not significantly altered by INTS11 knockdown. Notably, all these genes have AsiSI sites near their TSS. In order to minimize the effect of the elongation defect observed in INTS11-depleted cells on our expression analysis, we confined our gene expression analysis to the first 9 kb of the gene bodies. Fold change enrichment analysis revealed a significant decrease in the expression of AsiSI target genes in siCtrl-transfected cells treated with OHT compared with siCtrl-transfected cells treated with ethanol. However, compared with OHT-treated control cells, OHT-treated INTS11-knockdown cells showed reduced transcriptional silencing (Fig. 5e). These results were validated by quantitative RT-PCR on selected target genes (Fig. 5f), which provided further confirmation of the role of INTS11 in DSB-induced gene silencing. In summary, these findings suggest that Integrator, by providing a fail-safe mechanism, possibly through termination of a fraction of RNAPII complexes, plays an important role in preventing formation of spurious transcripts at DSB sites.

## Discussion

In this study, we used a combination of high-resolution genome-wide analyses (TT-seq, ChIP-seq and DRB/TT-seq) to gain insight into the role of INTS11 in regulating transcription by RNAPII. Overall, we observed a global reduction in nascent RNA synthesis on gene bodies, suggesting a possible elongation defect in INTS11-depleted cells. Notably, the elongation defect was detectable in all protein-coding genes longer than 15 kb and was not restricted to a certain class of protein-coding genes. Several lines of evidence presented in our study suggest that the elongation defect in INTS11-knockdown cells is due to diminished RNAPII processivity rather than decreased elongation rate. Most strikingly, DRB/TT-seq data showed that the rate of transcription is unchanged in INTS11-knockdown cells compared with control cells. ChIP-seq analysis of steady-state occupancy of RNAPII also revealed a significant reduction of the signal across the gene body in the knockdown cells, providing more evidence of the role of Integrator on RNAPII processivity.

Remarkably, we found a transition zone, beginning at ~ 12 kb and ending at ~ 35 kb from the TSS, where INTS11-knockdown cells exhibited a dramatic and progressive reduction in nascent transcription. Beyond this zone, the nascent transcription in the knockdown cells stayed constantly low. Our data thus propose the presence of a novel regulatory step exerted by Integrator that occurs within a region of 12 kb to 35 kb from the TSS, ensuring that RNAPII reaches its maximum processivity for efficient transcription elongation. In this regard, studies on RNAPII kinetics on human MCF-7 breast cancer cells have shown that upon release into productive elongation, RNAPII gradually accelerates into the gene body up to ~ 10–15 kb from the TSS, at which point it transitions to a final state of maximum elongation speed(Danko et al. 2013). Based on our data, we think that INTS11 may facilitate the efficient transition of RNAPII from the accelerating state to the fully processive state. The decrease we observe beyond the transition zone (TSS + 35 kb) in INTS11-knockdown cells is likely due to there being fewer RNAPII molecules able to pass through the accelerating phase into full processivity. The loss of processive RNAPII molecules is then amplified over the gene length and is therefore more noticeable at long genes. This observation is reminiscent of the role of Spt5 elongation factor, which was shown to be required for RNAPII processivity within a narrow window 15–20 kb from the TSS in mouse embryonic fibroblast cells(Fitz et al. 2018). Integrator-bound PP2A, through its phosphatase activity, was shown to dephosphorylate SPT5(Huang et al. 2020). While it would be interesting to investigate the interplay between SPT5 and Integrator on regulating RNAPII processivity, Hu et al. found that the maintenance of RNAPII processivity by SPT5 is independent of its phosphorylation(Hu et al. 2021). Moreover, Integrator phosphatase module is shown to be functional even in the absence of the endonuclease(Stein et al. 2022). Therefore, we think the observed changes in the processivity of RNAPII upon INTS11 knockdown is less likely to be an indirect effect on SPT5 function.

Previous studies have shown quality control mechanisms exerted by Integrator acting at promoter-proximal regions. Lykk-Andersen et al. demonstrated that Integrator at promoter-proximal regions (within ~3 kb of TSS) provides a quality control mechanism for non-productive transcription through termination of RNAPII complexes that are unfavourably configured for transcriptional elongation(Lykke-Andersen et al. 2021). In another study, Beckedroff et al. showed that Integrator is associated with paused RNAPII and, through its endonucleolytic cleavage activity, induces premature termination of nascent transcripts in the proximity of the +1 nucleosome, thus mediating transcriptional elongation at human promoters(Beckedorff et al. 2020). Recent study by Stein et al. showed that rapid degradation of INTS11 increased RNAPII release into early gene bodies. However, the released RNAPII is deficient in elongation and only short genes are upregulated. Consistent with these findings, we found that INTS11 depletion results in accumulation of RNAPII near TSSs and an increase in nascent RNA production of short genes. Our study further expands the knowledge on the role of Integrator on transcription and proposes a new role for Integrator in the transition of RNAPII to its maximal elongation state in the 10–15 kb region, for which no mechanism has yet been proposed.

Given that our results are based on INTS11, the main RNA endonuclease subunit of the Integrator complex, it is most plausible that INTS11, through its endonuclease activity, stimulates the cleavage and termination of low-processive, incompetent RNAPII molecules and in this way facilitates efficient elongation. Further studies using a catalytically inactive mutant of INTS11 will be required to confirm this hypothesis.

We also observed a genome-wide increase in TSS-upstream antisense transcripts upon INTS11 knockdown. Furthermore, we found that INTS11 and CTD 1YP are co-enriched on promoter upstream regions, where antisense transcription occurs. CTD 1YP has been identified as a hallmark of antisense promoter transcription(Descostes et al. 2014) and was shown to control the termination of 5′ antisense transcripts(Shah et al. 2018). Our data thus support a role for Integrator, in association with CTD 1YP, in processing non-productive antisense transcripts and, thereby, terminating RNAPII molecules with inefficient elongation capacity.

Although our study defines a positive role for the Integrator complex in transcription regulation, which is in line with previous studies performed on human cells(Beckedorff et al. 2020), *Drosophila* Integrator complex was shown primarily to attenuate gene expression – mostly acting on genes that are involved in signal-responsive pathways(Elrod et al. 2019). Nevertheless, using AsiSI cells, we found that INTS11 is important for attenuating gene transcription at DSB sites. Such transcriptional repression is thought to be critical for preventing the generation of truncated RNA molecules with undesirable functions(Shanbhag et al. 2010; Pankotai et al. 2012; Awwad et al. 2017; Machour and Ayoub 2020). We therefore conclude that Integrator mainly functions as a positive regulator of transcription, but it can also locally attenuate the spurious transcription associated with DSBs in DNA.

Different subunits of the Integrator complex were previously shown to have roles in DNA damage response and repair(Li et al. 2009b; Skaar et al. 2009; Zhang et al. 2009; Zhang et al. 2013). However, it is not well known whether Integrator functions as a whole complex or whether sub-modules also have distinct roles. Our study is the first report linking Integrator catalytic subunit INTS11 to transcriptional silencing at damage sites. Due to the lack of suitable INTS11 antibody for immunofluorescence assay, we were not able to examine INTS11 foci formation in AsiSI cells. Nevertheless, we showed the recruitment of another Integrator subunit, INTS12, to DSB sites in an RNAPII-dependent manner. Therefore, it is likely that INTS12 cooperates with INTS11 (and possibly with other subunits of the complex) in modulating the expression of AsiSI-target genes. In this regard, we found co-localization of INTS12 with CTD Y1P at the damage sites. CTD Y1P was shown to be required for transcription termination at the 5′ and 3′ end of genes and to be essential for recruitment of the Integrator complex(Shah et al. 2018). Taken together, these results suggest a strong functional link between CTD Y1P and Integrator, and they support a model where Integrator is recruited to the damage sites in a CTD-dependent manner to cleave the 3′ end of nascent RNA, causing termination of a fraction of genomic transcription that would otherwise be truncated due to the presence of DSBs.

Since our study was based on AsiSI-induced DSBs that mostly happen near the TSS, our analysis is restricted to gene expression at the 5′ end of the genes. Therefore, it would be interesting to further investigate the transcription-dependent role of Integrator on other genomic sites upon induction of DSBs.

In summary, our results add additional complexity to the role of Integrator in regulating RNAPII activity and propose the concept that Integrator, besides having a global positive role in transcription elongation, may target a subset of genes upon environmental stimuli to potentially repress their activity.

## Materials and Methods

### Cell culture and cell transfection

U2OS and AsiSI–ER–U2OS cells were cultured in DMEM (Thermo Fisher Scientific) supplemented with 10% FBS (Sigma-Aldrich) at 37°C in 5% CO_2_ incubator.

For siRNA transfection, cells were reverse transfected with Lipofectamine RNAiMAX (Thermo Fisher Scientific) reagent according to the manufacturer’s instruction. Human Silencer™ Select siRNAs against, INTS9 (#s31431), INTS11 (#s29893, #s29894, #s29895), and INTS12 (#s32713) were purchased from Thermo Fisher Scientific. Silencer™ Select Negative Control No. 1 siRNA (Thermo Fisher Scientific, #4390843) was used as a non-targeting siRNA (siCtrl). All siRNAs were used at 5 nM final concentration during transfection. All the experiments were done 48 hours post transfection. All the experiments were performed using siINTS11 #1 (#s29893) except the experiment in FigS1 which was performed using siINTS11 #2 and #3 (#s29894 and #s2989 respectively).

For AsiSI induction, AsiSI–ER–U2OS cells were incubated with 300 nM OHT in culture medium for different times.

### Immunoblotting

Cells were lysed in an appropriate volume of lysis buffer containing 20 mM HEPES pH 7.5, 0.5 M NaCl, 5 mM EDTA, 10% Glycerol and 1% Triton-X 100, containing protease inhibitor (Roche) and phosphatase inhibitor (Merck). After 10 cycles of sonication using Bioruptor Pico (Diagenode), supernatants were cleared of debris by centrifugation at maximum speed at 4°C.

Protein was quantified using the BCA assay (Thermo Fisher Scientific). Equal amounts of protein extract (20–30 *μ*g) were resolved by SDS-PAGE (4–15% gradient precast TGX polyacrylamide gel, Bio-Rad Laboratories), followed by standard western blot procedure. The following antibodies were used: anti-INT11 (Bethyl Laboratories, #A301-274A), anti-INTS9 (Cell Signaling, #13945) and anti-GAPDH (Abcam, #ab9485). Anti-mouse and anti-rabbit HRP-conjugated secondary antibodies were purchased from Cell Signaling. Immunoblots were developed with ECL reagents (Amersham) on Amersham Imager 680.

### Quantitative RT-PCR

Total RNA was extracted using miRNeasy kit (QIAGEN) with on-column DNase digestion. Extracted and purified RNA was then converted to cDNA using PrimeScript cDNA synthesis kit (TaKaRa) according to the manufacture’s instruction. Synthesized cDNA was used for subsequent real-time RT-PCR. Quantitative RT-PCR reaction was performed using SYBR PCR master mix (TaKaTa) in a QuantStudio 6 Flex Real-Time PCR System. Results were normalized to RPLPO expression and were plotted relative to control cells. Data are presented as mean ± S.D.

### Immunofluorescence

Immunofluorescence was performed on cells grown on coverslips. Cells were first fixed with 4% paraformaldehyde in PBS for 10 min and permeabilized with 0.2% Triton X-100 for 10 min at room temperature. Coverslips were blocked in PBS + 0.5% BSA for 20 min and incubated with primary antibody for 2 hours. Cells were washed with PBS and then incubated with secondary antibodies (Thermo Fisher Scientific). After 3 washes with PBS, coverslips were stained with DAPI (1:5000, Thermo Fisher Scientific) for 5 min. Slides were then mounted using Mowiol (Sigma-Aldrich). Slides were imaged using Zeiss Apotome.2 microscope and analysed with Fiji software. The following primary antibodies were used for immunofluorescence: RNAPII CTD phospho-Tyr antibody (Active Motif, #61383), Phospho-Histone H2A.X (Ser139) antibody (BioLegend, #613401 and Cell Signaling, #9718T), INTS12 antibody (Merk, #HPA035772).

### TT-seq

TT-seq was performed as previously described(Gregersen et al. 2020). Briefly, 48h post siRNA transfection, 4sU (1 mM final concentration) (Sigma-Aldrich) was added to the medium of the cells for 15 min. The medium was then discarded, and total RNA was extracted using TRIzol according to the manufacturer’s instruction (Thermo Fisher Scientific). Extracted RNA was then treated with DNaseI (Thermo Fisher Scientific) and purified using phenol/chloroform/isoamyl alcohol and ethanol precipitation. Around 100 μg purified RNA was spiked with 1 μg 4-thiouracil (4TU) labelled yeast RNA. The RNA mixture was fragmented to approximately 100-300 nucleotide fragments by sodium hydroxide treatment (0.16 M for 30 min on ice). The fragmentation step was then stopped by adding Tris pH 6.8 (0.4 M final concentration) and then samples were cleaned-up on Micro Bio-Spin P-30 gel columns (Bio-Rad Laboratories). 4sU-RNA was then biotinylated using 0.1 mg/ml MTSEA biotin-XX linker (Biotium, #BT90066) for 30 min at room temperature in the dark with rotation. Biotinylated RNA was purified by phenol/chloroform/isoamyl alcohol and ethanol precipitation. Purified biotinylated RNA was captured by μMACS streptavidin MicroBeads (Miltenyi Biotec) and after two washes was eluted with 100 μl of freshly prepared 100 mM DTT and then purified using RNeasy MinElute Cleanup kit (QIAGEN). 4sU-RNA was quantified with the Quit RNA High Sensitivity kit (Thermo Fisher Scientific). 500 ng RNA was used for library preparation using NEBNext Ultra II Directional RNA Library Prep kit (NEW ENGLAND BioLabs) according to the manufacturer’s protocol. Three independent replicates were performed for each condition (siCtrl and siINTS11). The libraries were pooled and sequenced on the HiSeq4000 (Illumina) using SE100 mode.

### DRB/TT-seq

Forty-eight hours after siRNA transfection, 5,6-dichloro-1-beta-D-ribofuranosylbenzimidazole (DRB) (Sigma-Aldrich) was added to the medium of the cells at final concentration of 100 μM. After 3 hours, cells were washed three times with PBS equilibrated at 37°C, and pre-warmed fresh medium was then added to the cells. 4sU (1 mM final concentration) was added to the medium during the last 10 min before cell harvesting. 10-, 20- and 30-min post DRB release, cells were harvested directly on the plate with TRIzol reagent (Thermo Fisher Scientific). 4sU-RNA pulldown was done as described for TT-seq. Two independent replicates were performed for each condition (siCtrl and siINTS11) and for each time (10-, 20- and 30-min post DRB release).

### TT-seq and DRB/TT-seq data analysis

First, the quality of sequencing was checked using FastQC. Adapters were trimmed from the 3’ ends of the reads using Trimmomatic v0.36(Bolger et al. 2014). Reads mapping to rRNA genes were removed using BBDuk in the BBTools package v36.20 [sourceforge.net/projects/bbmap]. The rRNA cleaned reads were then mapped to the human reference genome (hg38) with STAR aligner v2.7(Dobin et al. 2013). Putative PCR duplicates were marked and removed using MarkDuplicates (Picard tools v2.23.8) [https://broadinstitute.github.io/picard/].

Scale factor for each sample was computed by passing yeast spike-in gene count matrix to the estimateSizeFactors function in the Bioconductor DESeq2 package(Love et al. 2014). The aligned read files (BAM files) were sorted according to the chromosome using SAMtools v1.8(Li et al. 2009a). BAM files were converted to normalized strand-specific coverage files (bigwig files), which were used for downstream analyses, using bamCoverage, deepTools(Ramirez et al. 2016). Heatmaps and metaplots were generated with computeMatrix using deepTools. Tracks were visualized on the UCSC Genome Browser.

For differential gene expression analysis, reads were quantified using featureCounts from Subread package v2.0.3(Liao et al. 2014). Differentially expressed genes were obtained by DESeq2 tool on the gene count matrix using calculated yeast spike-in scale factors. Differentially expressed genes were defined as P-adjusted value of < 0.05 and a log_2_ fold change greater than 0 in either direction. Genes were considered as expressed if their RPKM (reads per kilobase per million sequenced reads) was greater than 0.5 (RPKM > 0.5).

Elongation rate was calculated by first fitting a smoothing spline to each DRB meta-profile (10, 20 and 30 min) and then finding the maximum value (wave peak) of the fitted spline curve at each time point as described in Gregersen et al. 2020(Gregersen et al. 2020).

### Total RNA-seq and data analysis

Total RNA was extracted using TRIzol reagent (Thermo Fisher Scientific). Genomic DNA was removed by Turbo DNase treatment (Thermo Fisher Scientific). Ribosomal RNAs were depleted using Illumina Ribo-Zerp Plus rRNA Depletion kit (Illumina). Ribosomal-depleted RNA was then used for library preparation using NEBNext Ultra II Directional RNA Library Prep kit (NEW ENGLAND BioLabs) according to the manufacturer’s protocol. The experiment was performed with two independent replicates for each condition (siCtrl and siINTS11). The libraries were pooled and sequenced on the HiSeq4000 (Illumina) in the 100-nt paired-end mode.

Raw FASTQ data were processed with Trimmomatic v0.36 to remove 3’ end adapters and low-quality reads and then aligned to the human genome hg38 using STAR aligner v2.7. After removal of PCR duplicates using MarkDuplicates (Picard tools v2.23.8), counts on exonic regions were extracted using featureCounts and RPKM was calculated for each gene.

Splicing efficiency was quantified using SPLICE-q with default parameters(de Melo Costa et al. 2021).

### ChIP-seq and data analysis

Cells grown in tissue culture dishes were first washed 2 times with PBS and then cross-linked with freshly prepared formaldehyde solution (final concentration 1%) for 10 min at room temperature. Fixation was quenched by adding Glycin (final concentration 0.25 M) for 5 min at room temperature. Cells were then washed 5 times with ice-cold PBS and harvested in PBS. After centrifugation, cell pellet was resuspended in ice-cold LB1 buffer (50 mM HEPES pH 7.5, 140 mM NaCl, 1 mM EDTA, 10% Glycerol, 0.5% NP-40, 0.25% Triton X-100) supplemented with protease and phosphatase inhibitors for 10 min on ice. After centrifugation, nuclei were extracted by resuspending the pellet in cold LB2 buffer (10 mM Tris-HCl pH 8.0, 200 mM NaCl, 1 mM EDTA, 0.5 mM EGTA) supplemented with protease and phosphatase inhibitors for 10 min on ice. Finally, the pellet was resuspended in cold LB3 buffer (10 mM Tris-HCl pH 8.0, 100 mM NaCl, 1 mM EDTA, 0.5 mM EGTA, 0.1% Na-Deoxycholate, 0.5% N-lauroylsarcosine) supplemented with protease and phosphatase inhibitors. Samples were then sonicated using Bioruptor Pico (Diagenode) to obtain DNA fragment size distribution of 150-300 bp.

For immunoprecipitation, cleared lysates were incubated with 10 μg of total RNAPII antibody (Bethyl Labratories, #A304-405A) previously bound to protein A and G Dynabeads (Thermo Fisher Scientific) for overnight at 4°C with rotation. Then, the beads were captured using DynaMag magnet (Thermo Fisher Scientific) and washed 6 times with ice-cold RIPA buffer (50 mM HEPES pH 7.5, 500 mM LiCl, 1 mM EDTA, 1% NP-40, 0.7% Na-Deoxycholate) and once with TE 1X+50 mM NaCl. Chromatin-protein complexes were eluted from the beads with elution buffer (1% SDS + 100 mM NaHCO_3_) for 15 min at 30°C with agitation. The beads were captured, and the supernatant was transferred to a new tube and treated with RNaseA (Thermo Fisher Scientific) at 65°C overnight followed by proteinase K digestion (Sigma Aldrich) at 60°C for 1 hour on a shaker. The DNA was purified using QIAquick PCR Purification kit (QIAGEN) and then quantified using Qubit dsDNA HS Assay kit (Thermo Fisher Scientific). Library preparation was done using NEBNext Ultra II DNA Library Prep Kit for Illumina (NEW ENGLAND BioLabs) as described by the manufacturer. The ChIP-seq experiment was done with two independent replicates for each condition (siCtrl and siINTS11). The libraries were pooled and sequenced on the HiSeq4000 (Illumina) using SE100 mode.

Adapters were trimmed using Trimmomatic v0.36. Obtained sequences were mapped to the human hg38 reference genome with Bowtie2. PCR duplicates were removed with Picard MarkDuplicates tools. Normalized bigwig files were generated for each bam file with bamCoverage tool (deepTools) using −normalizeUsing parameter. The normalization factor was calculated from the average counts of total RNAPII signal between siCtrl and siINTS11 samples across the gene bodies of a set of genes found to be non-affected in bulk RNA-seq data. Normalized bigwig files were used for generating meta-profiles.

### ChIP-SICAP-MS

AsiSI–ER–U2OS cells were treated with OHT (300 nM) for 1 hour. Cells were then cross-linked with 1.5% formaldehyde and processed for ChIP-SICAP as previously described(Rafiee et al. 2020; Rafiee and Krijgsveld 2021).

Raw MS data were analysed using MaxQuant(Cox and Mann 2008) (2.0.1.0) by default settings. In brief, MSMS spectra were searched against the UniProt (Swissprot) database (*Homo sapiens*) and database of contaminants. Trypsin/P and LysC were chosen as enzyme specificity, allowing a maximum of two missed cleavages. Cysteine carbamidomethylation was chosen as the fixed modification, and methionine oxidation and protein N-terminal acetylation were used as variable modifications. Global false discovery rate for both protein and peptides were set to 1%. The match-from-and-to, re-quantify, and intensity-based absolute quantification (iBAQ) options were enabled. The protein groups were processed in RStudio using R version 4.0.0. The proteins only identified by site, Reverse, and potential contaminants were filtered out. The limma package was used to determine Bayesian moderated t-test p-values and Benjamini-Hochberg (BH) adjusted p-values (adj. p-value or FDRs). We considered log2(FC)> 1 and and adj. p-value <0.05 as significantly enriched proteins.

**Figure 6.**
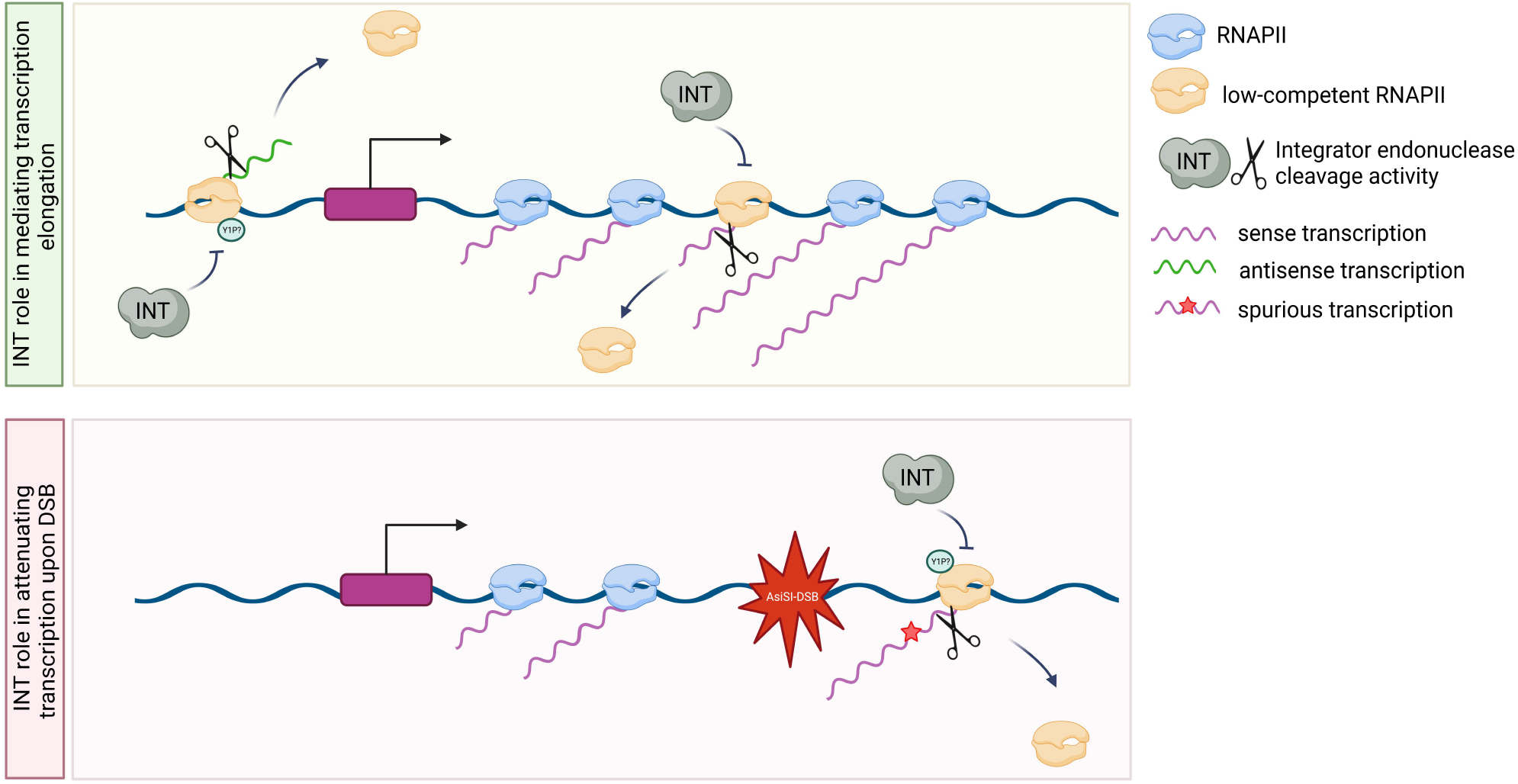
Model showing the role of Integrator in mediating transcription elongation in unperturbed condition (top) and in attenuating transcription upon formation of AsiSI-induced DSB (bottom).

## Data and code availability

All raw and processed data (TT-seq, DRB/TT-seq, ChIP-seq) are deposited in GEO under the accession number GSE210880. INTS11 ChIP-seq was downloaded from GEO accession GSE85098. mNET-seq data were downloaded from GEO accession GSE81662.

The mass spectrometry proteomics data have been deposited to the ProteomeXchange Consortium via the PRIDE(Perez-Riverol et al. 2022) partner repository with the dataset identifier PXD037045.

Only publicly available tools were used in data analysis as described in the Materials and Methods section.

## Acknowledgments

We thank Robert Goldstone and Deb Jackson from the Crick Advanced Sequencing Facility for excellent technical support in NGS sequencing. Cartoons in Fig 3h and Fig 6 were created with BioRender.com. This work was supported by the Francis Crick Institute which receives its core funding from Cancer Research UK (FC010110; 215593/Z/19/Z), the UK Medical Research Council (FC010110), and the Wellcome Trust (FC010110). For the purpose of Open Access, the author has applied a CC BY public copyright licence to any Author Accepted Manuscript version arising from this submission. N.M.L. and J.U. are additionally funded by a Wellcome Trust Joint Investigator Award (103760/Z/14/Z). N.M.L receives core funding from the Okinawa Institute of Science & Technology Graduate University.

## Author contributions

S.R. conceived the study, designed and performed most of the experiments and the bioinformatics analyses. M.R. designed and performed mass spectrometry analysis. S.R. discussed and interpreted the results with inputs from M.R., N.M.L. and J.U. S.R. wrote the paper with input from the other authors.

